# Genetic analysis of *rab7* mutants in zebrafish

**DOI:** 10.1101/2023.03.09.531857

**Authors:** Daniel Heutschi, Etienne Schmelzer, Vahap Aydogan, Alexander Schmidt, Heinz-Georg Belting, Anne Spang, Markus Affolter, Maria P. Kotini

## Abstract

Vascular network formation requires the fusion of newly formed blood vessels and the emergence of a patent lumen between the newly established connections so that blood flow can start. Lumen formation has been shown to depend on the late endosomal/lysosomal pathway in various organs of animal tubular systems. Here, we identified a late endosomal/lysosomal vesicular fraction (Rab7/Lamp2) in early zebrafish angiogenic sprouts, which appears to contribute to apical membrane growth during lumen formation. To study the effect of the late endocytic pathway on vascular development, we generated mutant alleles for all three *rab7* genes in zebrafish (*rab7a, rab7ba, rab7bb*). All *rab7* genes are expressed in wild-type zebrafish and we did not detect any compensatory effects by the other *rab7* isoforms in single knockout mutants, which were all viable. Only the triple mutant was lethal suggesting some functional redundancy. However, the different *rab7* isoforms fulfil also at least partially independent functions because eggs laid from mothers lacking two *rab7* (*rab7a and/or rab7bb*). showed reduced survival and contained enlarged yolk granules, suggesting maternal contribution of these two *rab7*. Finally, we observed minor effects on lumen formation in embryos which still express one copy of *rab7*. Our results support the notion that the late endocytic/lysosomal compartment contributes to lumen expansion.

## Introduction

The vasculature is the first organ to form in the vertebrate embryo. Its function is to supply the surrounding tissue with nutrients and oxygen as well as immune cells and it is vital for the growth of the embryo. An essential step in the establishment of a functional network is the formation and expansion of the vascular lumen. The onset of lumen formation begins with the apical polarization of endothelial cells (ECs) upon contact of two vessel segments (anastomosis) and the expansion of the initial apical patch in a disc-like structure. In the final step, the lumens of the two contacting vascular branches must be connected to allow for blood flow. Lumen expansion has been described to occur via two distinct processes, transcellular lumen formation and cell rearrangements, a process also referred to as cord hollowing (Ellertsdóttir et al., 2010; Herwig et al., 2011).

During cord hollowing, the lumen is formed between ECs, while during transcellular lumen formation, the lumen is formed within ECs and is driven by blood pressure. The transcellular lumen forms via membrane invagination through the cell body and subsequent fusion of the invaginating apical membrane with the newly-formed, distal apical patch. The process then progresses into the next cell (Francis et al., 2022; Gebala et al., 2016; Herwig et al., 2011; Lenard et al., 2013). It is not clear which membranous cell compartment contributes to the formation and the enlargement of the apical membrane.

The Rab GTPase vesicle trafficking program, and more specifically Rab35, has recently been shown to regulate the establishment of apicobasal polarity during angiogenesis *in vitro* and *in vivo* in the zebrafish embryo (Francis et al., 2022). Apart from the vasculature, vesicular trafficking has been involved in the formation of lumen in other systems.

The mechanism of lumen connection in the vertebrate vasculature shares many morphological similarities with the one in the tracheal system of *D. melanogaster* (Camelo et al., 2022; Caviglia and Luschnig, 2014; Hayashi and Kondo, 2018; Kotini et al., 2019). In the embryonic trachea, the formation of a continuous lumen via apical membrane fusion has been described to be dependent on vesicular trafficking (Caviglia et al., 2016). In the absence of the tethering protein Unc-13-4/Staccato, individual branches of the tracheal system fail to connect their lumens and do not form a continuous network. In tracheal fusion cells, Unc-13-4/Staccato recruits vesicles that have been characterized by the presence of Rab7, Rab39 and Lamp1 as secretory lysosomes, which accumulate in the cytoplasm between the two growing luminal membranes of the fusion cell (Caviglia et al., 2016).

Formation of late endosomes and lysosomes is dependent on Rab7. In the endocytic pathway, Rab7 is recruited to early endosomes, converts them into late endosomes and promotes their fusion with lysosomes (Marwaha et al., 2017; Poteryaev et al., 2007, 2010; Rojas et al., 2008). Rab7 recruits the HOPS tethering complex that interacts with SNARE proteins and leads to membrane-membrane recognition and mediates fusion of late endosomes with lysosomes (Bröcker et al., 2012; Solinger and Spang, 2013). Loss of Rab7 leads to severe defects in early development. In *C. elegans,* yolk granules are enlarged upon reduction of Rab7 by RNAi or upon a knockdown of its guanine exchange factor (GEF) SAND-1; loss of Rab7 causes embryonic lethality (Poteryaev et al., 2007). In mice, the absence of Rab7 yield a loss of endoderm specification due to a lack of Wnt signalling (Kawamura et al., 2012).

Since the precise role of late endocytic trafficking in vertebrate vascular lumen formation is not known, and since Rab7 is a main organizer of late endosomal trafficking, we analysed the expression of EGFP-Rab7a during lumen formation/expansion in zebrafish embryos. We find that Rab7a colocalizes with dot-like structures also marked by a CAAX membrane marker. These structures often elongate along the apical membrane, suggesting that they fuse with the latter. The dot-like structures also colocalize with Lamp2, a lysosomal-associated membrane protein. These results suggest that a late endosomal, lysosomal compartment might contribute to apical membrane growth in angiogenesis.

To analyse the role of Rab7 in vascular development, we generated mutant alleles for the three *rab7* genes in zebrafish, *rab7a*, *rab7ba* and the newly found *rab7bb*, which we analysed in this study. We found that this third *rab7* gene, *rab7bb*, shares some redundant function with *rab7a.* We also found that loss of maternally contributed *rab7* leads to an increase in yolk granules, similar to what was observed in *C. elegans*, and that complete loss of Rab7 in triple mutants is lethal. High resolution confocal imaging revealed that lumen formation in the analysed double mutants is not significantly impaired. In order to study the role of Rab7 in angiogenesis and lumen formation, endothelial-specific knockout or knockdown strategies will be needed.

## Results

### Rab7 colocalises with mCherry-CAAX at dots

To visualize membrane dynamics during vascular lumen formation, we used a transgenic reporter *Tg(kdrl:HsHRAS-mCherry)^s916^*, which effectively labels membrane compartments in endothelial cells. Previous live-imaging studies have shown that this reporter preferentially labels the apical membrane of nascent blood vessels (Lenard et al., 2013; Phng et al., 2015). We therefore reasoned that this reporter is ideally suited to detect and identify vesicular membrane structures which contribute to the nascent apical cell membrane. Time-lapse analysis revealed local accumulation of mCherry-CAAX protein in the cytoplasm as dots (Fig 1A-B).

**Figure 1.**
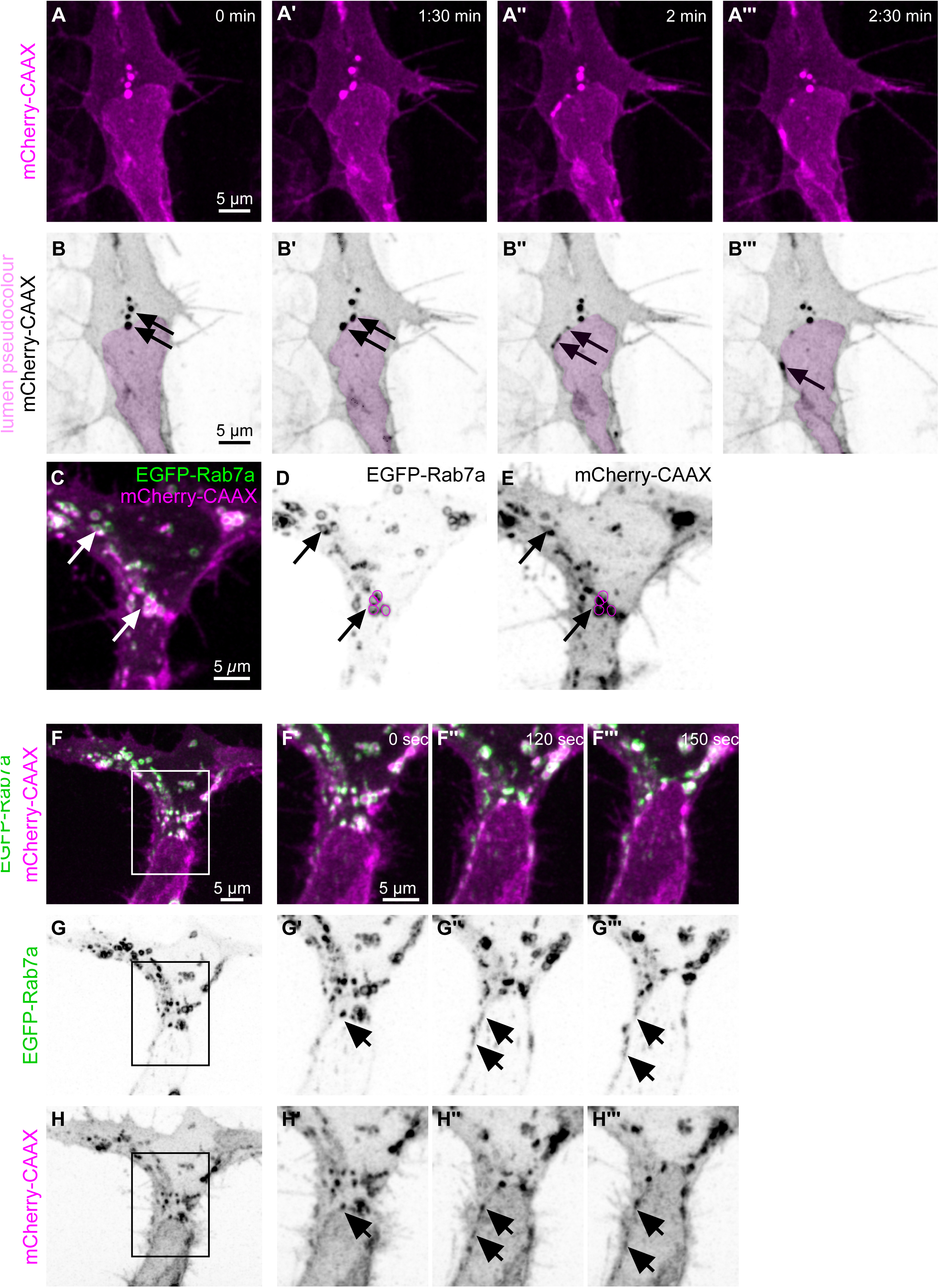
Rab7 co-localization with the apical marker CAAX and elongation of CAAX/Rab7 dots at the apical membrane. **A-B** Confocal images from a time-lapse of a tip cell from a transgenic *Tg(kdrl:mCherry-CAAX)^S916^* embryo shown in magenta. B-B’’’ Inverted contrast of mCherry-CAAX from A shows CAAX dots which elongate along the apical membrane. Pink pseudo-color indicates the vascular lumen. **C-E** Confocal images of the tip cell of a double transgenic *Tg(fli:eGFP-Rab7a)^ubs48^; Tg(kdrl:mCherry-CAAX)^S916^* embryo at 32hpf . **D** Inverted contrast image of the EGFP-Rab7 channel. **E** Inverted contrast image showing the membrane marker mCherry-CAAX in ECs. Arrows point to co-localisation of Rab7 and CAAX at dots. **F** Distribution of Pearson Correlation Coefficient (PCC) of EGFP-Rab7a ROIs in correlation to mCherry-CAAX, showing a correlation between EGFP and mCherry signal (n=8 tip cells, N>3 different embryos). **F** A tip cell from a transgenic *Tg (kdrl:mCherry-CAAX)^S916^* embryo at 30hpf, transiently expressing *fli:EGFP-Rab7a*. **F’-F’’’** Stills from the ROI from D showing EGFP-Rab7 and mCherry-CAAX dots that move along the expanding apical membrane. **G-G’’’** EGFP-Rab7a signal alone. **H-H’’’** mCherry-CAAX signal alone. Arrows point to the EGFP-Rab7a and mCherry-CAAX dots.

In order to identify the nature of these CAAX dots, we transiently expressed different EGFP-fusions of Rab proteins, such as Rab5c (marker for early endosomes), Rab7a (marker for late endosomes) and Rab11a (marker for recycling endosomes) in the developing vasculature. No clear colocalization was observed with Rab5c and Rab11 (Fig S1). While mCherry-CAAX appeared as filled dots, EGFP-Rab7a was visible in doughnut-like structures around these dots (see arrows in Fig 1C). The colocalization of mCherry-CAAX and Rab7a suggests that these structures represent late endosomal-lysosomal compartments, and that such a compartment could contribute to apical membrane growth in angiogenic sprouts. Indeed, when we compared EGFP-Rab7a and mCherry-CAAX localisation during lumen expansion, we observed co-migration of Rab7/CAAX structures along the apical membrane (Figure 1G-I), suggesting that they might integrate into the apical membrane and may be a major source for the growth of this membrane compartment.

### Rab7a colocalizes with Lamp2 in endothelial cells

To further confirm the nature of the mCherry-CAAX/GFP-Rab7a-labelled structures as late endosomal/lysosomal, we made use of a BAC transgenic zebrafish line expressing a Lamp2-RFP fusion protein (Rodríguez-Fraticelli et al., 2015). Lamp2 is a component of late endosomal/lysosomal compartments, and is expressed in many different tissues in early zebrafish embryos, including the vasculature (see Fig 2A and C). Transient expression of EGFP-Rab7a in the vasculature of an embryo expressing Lamp2-RFP showed that Lamp2-RFP formed dot-like and doughnut-like structures in ECs, and that these structures indeed co-localized with EGFP-Rab7a (Fig 2A-2C and higher magnification in Fig 2A’-2C’, respectively). Strikingly, we observed that Lamp2-RFP-positive structures elongated along the apical membrane, similar to what we have seen previously for mCherry-CAAX dots (Fig 2D-2F; see also Fig 1A-B). We therefore conclude that mCherry-CAAX-positive structures represent late endosomal/lysosomal compartments, and that these compartments might contribute to the growing apical membrane during transcellular lumen formation.

**Figure 2.**
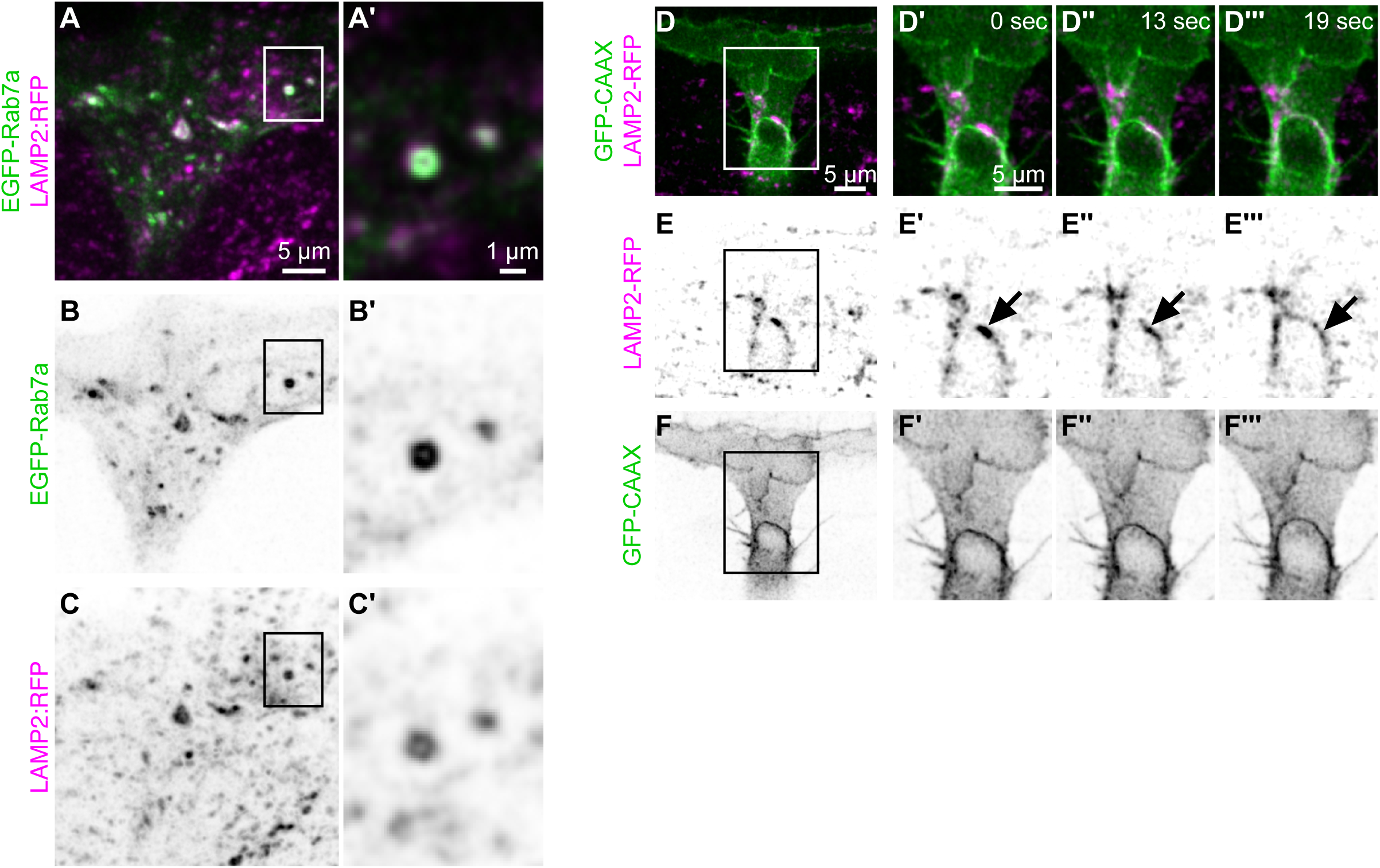
EGFP-Rab7a and Lamp2-RFP colocalize at dots. **A-B** Confocal images of a tip cell from a transgenic *TgBAC(Lamp2-RFP)^pd1117^* embryo at 30hpf, transiently expressing *fli:GFP-Rab7a*. **A** Z-projection of the tip cell. **A’** Single z-slice of the ROI in A. Two vesicles are depicted with signal positive for GFP-Rab7a and Lamp2-RFP. The largest vesicle shows a reduced signal in the centre for both GFP-Rab7a and Lamp2-RFP, resembling late endosomal/lysosomal structures. **B-B’** EGFP-Rab7a signal alone in endothelial cells. **C-C’** Lamp2-RFP signal alone. **D** Schematic representation of the blood vessel (tip EC) undergoing transcellular lumen formation. Rab7, Lamp2 and CAAX co-localise at dots which migrate along the expanding apical membrane. **D** Tip cell of a double transgenic *Tg(kdrl:GFP-CAAX)^s916^; TgBAC(Lamp2-RFP)^pd1117^* embryo at 34hpf. **D’-D’’’** Timelapse from the ROI in E showing Lamp2-RFP dot-like structure elongating along the apical, invaginating membrane. **E-E’’’** Lamp2 signal alone. **F-F’’’** mCherry-CAAX signal alone. Arrows point to the EGFP-Rab7 and mCherry-CAAX dot-like structures

### Identification and description of rab7 genes in zebrafish

As a first step to generate mutants for the *rab7* genes in zebrafish, we analysed and characterised all *rab7* genes encoded in the zebrafish (*Danio rerio*) genome. Two genes encoding Rab7 proteins have been described in zebrafish; *rab7a* and *rab7b* (Hall et al., 2017). A third gene (*zgc:100918)* has been proposed to encode a Rab7-like protein (Bayés et al., 2017). The amino acid sequences encoded by these three paralogues are 88% identical, with *zgc:100918* seemingly being a “hybrid” between *rab7a* and *rab7b*, showing 91% and 92% similarity to *rab7a* and *rab7b*, respectively (Fig S2A). A phylogenetic analysis of the amino acid sequences encoded by the three zebrafish *rab7* genes and *rab7* genes of mice and humans revealed three clusters. The first cluster is an interspecies *rab7a* cluster. The second cluster contains the *rab7b* genes from mice and humans and the third cluster consists of zebrafish *rab7b* and *zgc:100918*. This shows that *zgc:100918* is not a copy of *rab7a* but rather an ancient duplication of *rab7b* (Fig 3A). To further test this, we built the phylogenetic tree of the three *rab7* genes of different cyprinid species, which showed that the entire cluster of *zgc:100918* groups closer to the *rab7b* cluster than to *rab7a* cluster (Fig 3B). Investigation of the region around *zgc:100918* on chromosome 10 (Chr. 10) shows that genes in that region are annotated copies of genes within the region around *rab7b* on chromosome 8 (Chr. 8). In summary, sequence similarity, phylogenetic analysis and conserved synteny indicate that *zgc:100918* represents a second *rab7b* paralog, which we therefore name *rab7bb* hereafter, while *rab7b* will be referred to as *rab7ba*. According to two published transcriptomics databases (https://www.ebi.ac.uk/gxa/experiments/E-ERAD-475) (Lawson et al., 2020), *rab7bb* is expressed at similar levels as *rab7a* (Fig 3D), but at much higher levels than *rab7ba*, both in endothelial and non-endothelial cells (Fig 3D). Additionally, *rab7a* and *rab7bb* showed quite strong expression at the RNA level at the one cell stage, indicating maternal deposition of *rab7a* and *rab7bb,* but not of *rab7ba* (Fig 3E). Taken together, these data strongly suggest that the initially proposed *rab7*-like gene *zgc:100918* represents a copy of *rab7b*, and since its expression pattern is similar to *rab7a*, it has to be included in a genetic analysis of *rab7* in zebrafish.

**Figure 3.**
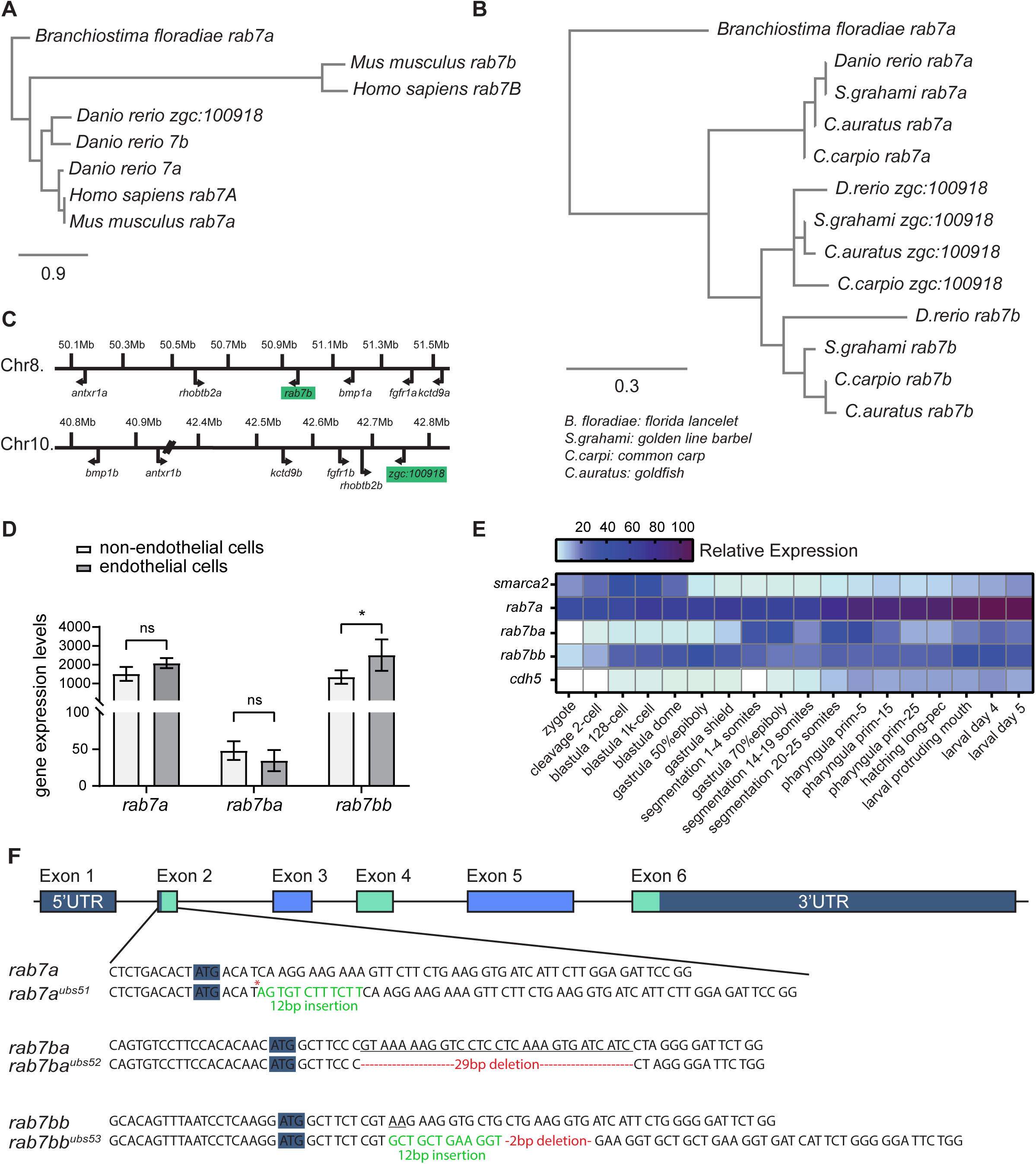
Description of all *rab7* isoforms in zebrafish. **A** Phylogenetic tree constructed from the protein sequence derived from amphioxus (outgroup), human, mouse and zebrafish *rab7* genes, scale: 0.9 amino acid substitutions. **B** Phylogenetic tree constructed from the protein sequences of *rab7a, rab7b* and *zgc:100918* genes from zebrafish, other cyprinid family members and amphioxus (outgroup), scale: 0.3 amino acid substitutions. **C** Representation of the chromosomal region around the genes *rab7b* on chromosome 8 and *zgc:100918* on chromosome 10. Indicated are the genes that are already annotated copies of each other. **D** Analysis of the expression of *rab7* genes in endothelial and non-endothelial cells from zebrafish transcriptomics data (Lawson, Li et al. 2020). **E** Heatmap of expression of *rab7* genes, a maternally contributed gene (*smarca2*) and an endothelial-specific gene (*cdh5*) during zebrafish development, based on data from an EMBL expression atlas (Papatheodorou et al. 2020 see Material and Methods). **F** Schematic representation of the gene structure of *rab7* genes in zebrafish. 5’ and 3’ UTRs and ATG (start codon) are highlighted in dark blue, alternating coding exons are represented in light green and light blue. Sequence of the exons 2 of *rab7a, rab7ba* and *rab7bb*. Below each gene sequence appears the respective sequence of mutant alleles *ubs51*, *ubs52* and *ubs53*. Deleted base pairs are underlined at the wild-type sequence, inserted base pairs are represented in light green and deletions are shown in red in mutant alleles. Red asterisk shows premature stop codon in the sequence of exon2.

### Characterization of rab7 protein levels in corresponding rab7 mutants

To investigate the role of *rab7* in vascular lumen formation, mutant alleles for all three *rab7* loci were generated using the CRISPR/Cas9 system. gRNAs were designed for each of the three *rab7* genes within the first coding exon (exon2) and as close as possible to the start codon to minimize the potential to generate a residual, functional protein. For *rab7a,* a 12 bp insertion leading to a stop codon after the second amino acid (aa) was isolated; this allele will be referred to as *rab7a^ubs51^*(Fig 3F; Fig S2D). For *rab7ba,* a 29 bp deletion leading to an out-of-frame protein after aa 4 and a stop codon in exon3 was isolated; this allele will be referred to as *rab7ba^ubs52^* (Fig 3F; Fig S2D’). Finally, for *rab7bb,* a 12 bp insertion accompanied by a 2 bp deletion leading to an out-of-frame protein after aa 4 and a stop codon in exon 3 of the gene was isolated; this allele will be referred to as *rab7bb^ubs53^* (Fig 3F; Fig S2D’’). Usage of unpredicted downstream alternative start codons in these mutants would lead to shortened Rab7 proteins lacking an important prenylation site (PS), which is encoded within the first 25 bp of the *rab7* genes (Fig 4A; Fig S2 A); without this site, the C-terminal XCXC domain cannot be prenylated and the protein cannot be inserted into the membrane (Sanford et al., 1995).

**Figure 4.**
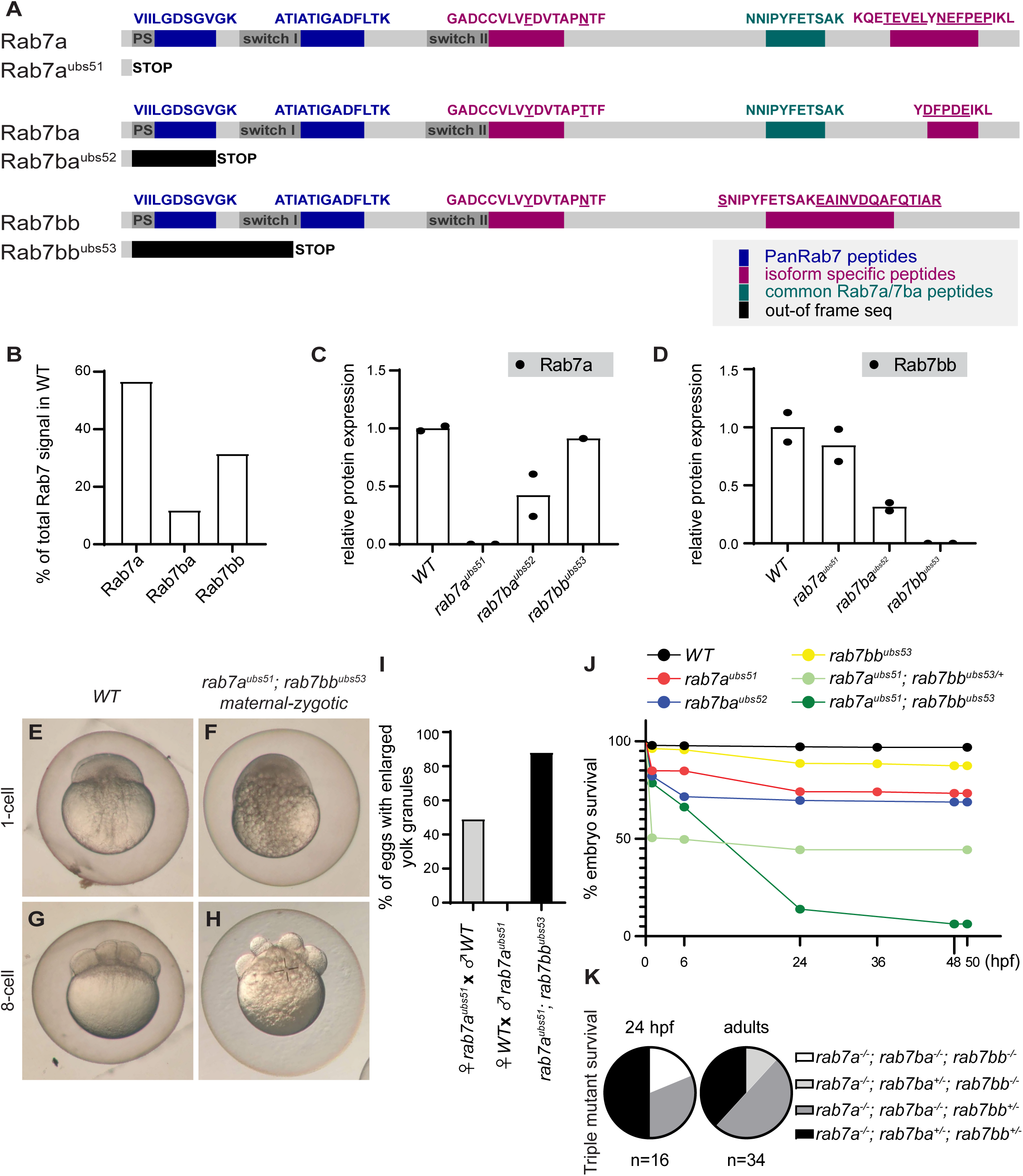
Characterisation of *rab7* mutant isoforms. **A** All three *rab7* amino acid sequences and their predicted mutant sequences. Peptides used for the Mass Spectrometry experiments are shown on top of each sequence. Peptides that recognise all three isoforms are shown in blue (PanRab7), isoform-specific peptides are shown in magenta and the common peptide for Rab7a and Rab7ba is shown in green. **B** Graph showing the contribution in % of the individual Rab7 isoforms to total Rab7 protein in wild-type (n= 2 pools of 20 embryos). **C and D** Individual value scatter plots of relative protein expression of the three different Rab7 isoforms. Levels were measured in two different pooled samples of wild-type, *rab7a mat-zyg, rab7ba mat-zyg* and *rab7bb mat-zyg* homozygous embryos. Values were then normalized to total amount of protein measured per sample and to the amount of wild-type sample (n= 2 pools of 20 embryos). **G-J** Bright field images of wild-type or maternal-zygotic *rab7a; rab7bb* double homozygous mutant embryos at 1-cell stage and 8-cell stage. **K** Bar graph showing the percentage of embryos with enlarged yolk granules in different *rab7* mutant crosses (n= 289-515 embryos, N=3 different single crosses per condition). **L** Survival plot of clutches from different mutant crosses. Percentage of surviving embryos from wild-type and *rab7a, rab7ba* and *rab7bb* homozygous incrosses, as well as incrosses from *rab7a* homozygous*, rab7bb* heterozygous adults and *rab7a; rab7bb* double homozygous parents (n=499-1438 embryos N= 2-7 crosses per condition.). **M** Triple mutant survival from a cross of a *rab7a^-/-^; rab7ba^-/-^; rab7bb^+/-^* mother and a *rab7a^-/-^; rab7ba^+/-^; rab7bb^-/-^* father (n=16 embryos at 24hpf, 34 adults after 3 months).

To test whether these mutant alleles represented indeed true null mutations (lacking specific Rab7 isoforms), targeted LC-MS proteomics analyses of homozygous in-crosses from each mutant allele were performed with pools of 24-hour old embryos. Since homozygous mutants of all three *rab7* mutant alleles were viable and fertile, we analysed the respective maternal-zygotic mutants for maternal contribution from each respective allele. Different peptides were used for the MS analyses, which were either common to all three proteins (PanRab7), to two of the three isoforms (Rab7a/Rab7ba), or specific for a given protein (see Fig. 4A and Methods). Care was taken to compare different protein isolates with the same peptide(s), rather than measuring the levels of different peptides using a single protein preparation. For a detailed description of the methods and the approaches taken, see the Method section.

In wild-type embryos, we found that the most abundant Rab7 protein was Rab7a (roughly 60% of the total amount of Rab7), while Rab7bb represented 30%. The least abundant isoform was Rab7ba, which represented roughly 10% (Fig 4B). The low levels of Rab7ba did not allow us to use the specific peptides in the mutant analyses (see below).

Our MS analyses using mutant embryos showed that *rab7a^ubs51^* and *rab7bb^ubs^* most likely represented null mutants, since no residual proteins or shortened fragments thereof were detected in the corresponding mutants (Fig 4C and Fig 4D, respectively). To measure potential residual protein levels of Rab7ba in *rab7ba^ubs52^,* a pan Rab7a-Rab7ba reference peptide was used (Fig 4A), because the reference peptides specific for Rab7ba (Fig 4A) were only detected at very low levels in the wild-type, but often remained under the detection threshold in the different mutants (see Method section for further details and explanations). Nonetheless, based on these analyses (Fig S2B), we conclude that *rab7ba^ubs52^*also represents a null allele.

The targeted LC-MS data further revealed that there is no obvious compensation; the levels of the remaining, wildtype Rab7 isoforms were not elevated in any of the *rab7* mutants we analysed; Rab7a was not elevated in *rab7ba^ubs52^* nor in *rab7bb^ubs53^*, while Rab7bb was not increased in *rab7a^ubs51^* nor in *rab7ba^ubs52^*(see Fig 4B, C). Again, we came to the same conclusion with respect to Rab7ba using a more indirect quantification approach, namely that its levels are not significantly increased in the absence of either of the two other isoforms (see Method section).

### Viability of rab7 mutant alleles

Since *rab7* is expected to play an important role in cell survival in general, we investigated the viability of the generated null mutant alleles. Single mutants of *rab7a, rab7ba* and *rab7bb* resulting from heterozygous in-crosses were viable and fertile (Fig S2E-F). Adults arising from these crosses showed normal mendelian distribution after 3 months (Fig S2G). Given the transcriptomics data, we argued that maternally deposited mRNAs or protein could rescue certain defects in homozygous embryos coming from heterozygous crosses. Using homozygous mutant parents for genetic crosses (thus removing potential contribution of maternally deposited mRNA and protein), we found a higher mortality in all *rab7* single mutants (Fig 4J). Survival rate drops from 97% in wild-type embryos to 84% in *rab7bb* homozygous mutants, to 59% for *rab7a* and to 76% in *rab7ba* homozygous mutants.

To investigate whether there was some degree of redundancy in function between the three *rab7* alleles, maternal-zygotic double mutants were analysed. While *rab7a*; *rab7ba* double homozygous fish did not show any change in viability compared to their single mutant variants, *rab7a*; *rab7bb* double homozygous mutants showed a drastic decrease in viability. The survival of offspring from in-crosses of *rab7a ^-/-^*; *rab7bb ^+/-^* fish was reduced to 40%. In double homozygous in-crosses, survival was reduced to 8% within the first 24 hours (Fig 4J). These results show that *rab7bb* shares some redundant function with *rab7a* and that the lowly-expressed *rab7ba* plays indeed a less important role for zebrafish embryo survival.

Despite the high lethality of *rab7a*; *rab7bb* double maternal-zygotic mutant embryos, roughly 10% of the embryos did survive. To test whether this might be due to residual *rab7ba* protein, we crossed fish that result in triple homozygous mutant progeny. We crossed a *rab7a^-/-^; rab7ba ^-/-^; rab7bb ^+/-^* female to a *rab7a^-/-^; rab7ba ^+/-^; rab7bb ^-/-^* male and screened the clutch for triple homozygous embryos using a four-primer multiplex PCR assay for each gene (for further details, see Methods). Out of 16 embryos, 3 were triple homozygous (Fig 4K), which is in accordance with the expected mendelian rate of 1/8. However, when screening 3 months old siblings of the same cross, 0/34 adult fish were triple homozygous. These data demonstrate that zebrafish which lack all Rab7 proteins are not viable.

### Increase of yolk granule number in rab7 maternal-zygotic homozygous mutants

Aside from embryo survival, loss of Rab7 may have severe defects in early development. A very early role linked to *rab7* function is the endocytic traffic pathway resulting in the formation of yolk granules. Loss of Rab7 or its GEF leads to a failure of yolk proteins to reach yolk granules in *C. elegans* (Poteryaev et al., 2007). In zebrafish, 1-cell-stage embryos of single *rab7a* or double *rab7a; rab7bb* mutants showed an increase in the size of yolk granules (Fig 4E-4I). This phenotype reflects a similar phenotype described in *C. elegans* (Poteryaev et al., 2007) and appears to be due to maternal contribution, since in progeny coming from *rab7a* homozygous mothers that were crossed to wild-type males, 49% of the embryos showed this phenotype. In contrast, 0% of embryos coming from wild-type mothers, crossed to *rab7a* homozygous males, showed any defects linked to yolk granules. The effect became even stronger in progeny from *rab7a; rab7bb* double homozygous adults. In this scenario, 88% of embryos showed an increase in yolk granules (Fig 4I). Coincidently, the frequency of occurrence of this phenotype was similar to the percentage of embryos that did not survive in these crosses. In fact, the majority of the 10% surviving embryos in these double homozygous in-crosses did not show this yolk phenotype; however, in rare cases, even embryos with yolk granules survived more than 24hpf. Occasionally, these embryos also showed defects in cell spacing in the blastodisc. In Fig 4H, the usually well-spaced organization of the cells (Fig 4F) was lost in eggs with yolk granules (Fig 4H).

### Lumen formation in rab7 single and double mutants is only slightly impaired

During blood vessel anastomosis, transcellular lumen formation occurs upon the formation of tip cell contacts and can be followed using the membrane marker *Tg(kdrl:mcherry-CAAX)* and a driver line forcing its expression in endothelial cells. The apical lumen front can be observed in trunk blood vessels roughly from 30 hpf onwards, expanding from the dorsal aorta dorsally through the newly forming vessels. Once the newly formed vascular loops have opened up to allow blood flow, blood pressure increases and the lumen diameter expands (Fig 5A). To investigate whether Rab7 plays a role in lumen formation in zebrafish as suggested by studies in other systems (Caviglia et al., 2016, 2017), *Tg(kdrl:mcherry-CAAX)* embryos were analysed by confocal live imaging in order to follow initial lumen formation, lumen fusion and lumen maintenance in *rab7* mutant fish. Lumen formation was observed in embryos homozygous mutant for each of the three *rab7* loss-of-function alleles and in embryos homozygous for the two *rab7* double mutants analysed (*rab7a; rab7ba* and *rab7a; rab7bb*) (Fig 5B-E); all blood vessels analysed formed a lumen during anastomosis and subsequently maintained it until the end of data acquisition. To quantify lumen maintenance, the diameter of the lumen was measured perpendicular to the vessel orientation at three different positions (Fig 5F). Lumen diameter was slightly reduced in *rab7ba* homozygous mutants as well as in *rab7a*; *rab7ba* double homozygous mutants (Fig 5G). Keeping in mind the strong maternal contribution of *rab7a* and *rab7bb*, we wanted to analyse whether this contribution plays a role during anastomosis and analysed embryos from homozygous in-crosses. These embryos showed a significantly reduced diameter in maternal-zygotic homozygous *rab7ba* mutants and maternal-zygotic homozygous *rab7a; rab7ba* double mutants. The lumen diameter was increased when comparing embryos that were zygotic homozygous mutants for *rab7a,* to maternal-zygotic homozygous mutants for *rab7a* or to double maternal-zygotic homozygous mutant for *rab7a; rab7bb*. Similar to the lethality studies, this indicates again that *rab7a* and *rab7bb* share some redundant function.

**Figure 5.**
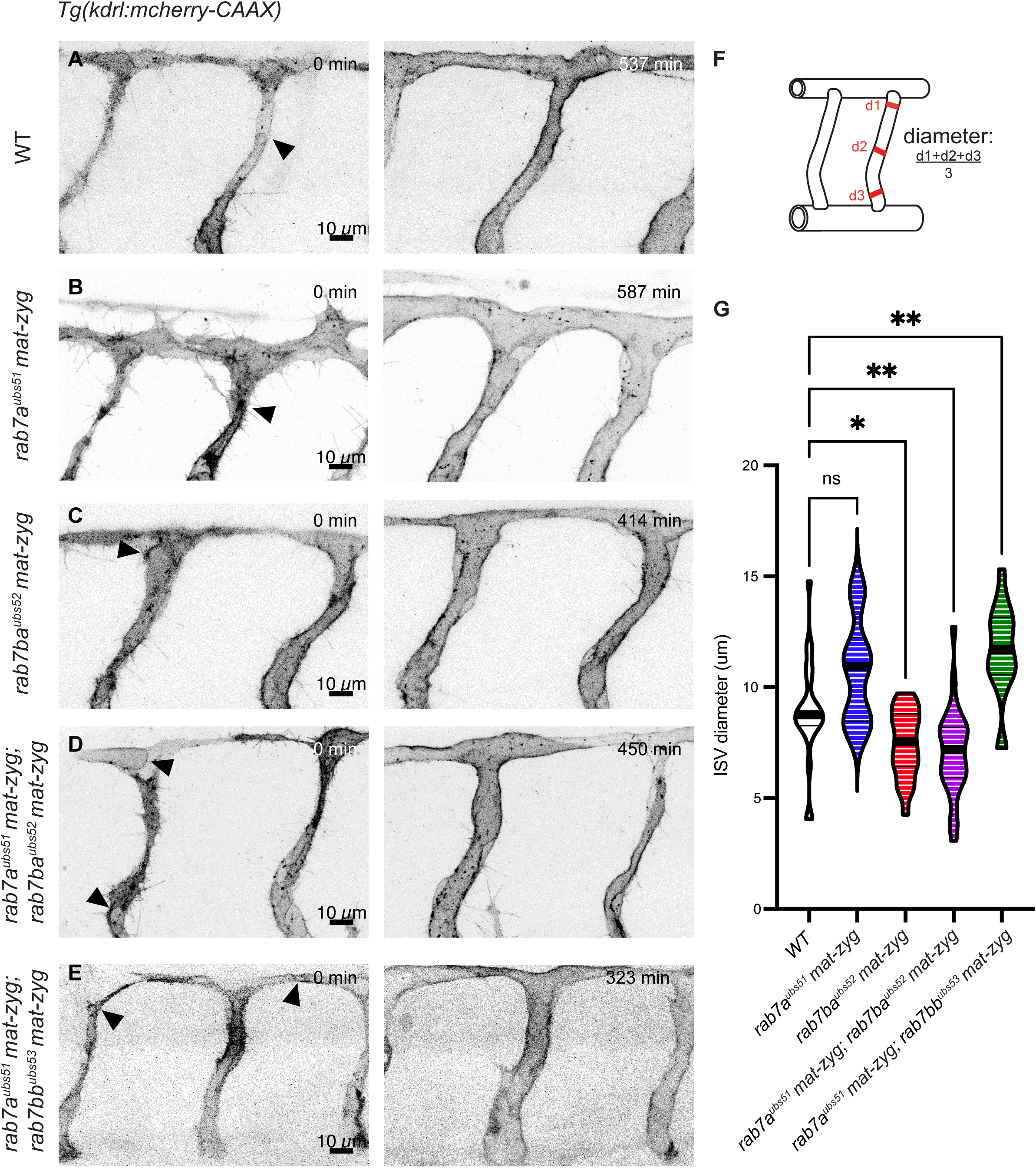
Vascular lumen defects in *rab7* mutants. **A-E** Confocal still pictures from time-lapse movies from transgenic *Tg(kdrl:EGFP-CAAX)^s916^* embryos at 34-44 hpf showing blood vessel lumenization in wild-type **(A),** maternal zygotic homozygous mutant for *rab7a* **(B)**, *rab7ba* **(C)**, maternal-zygotic double homozygous mutant for *rab7a; rab7ba* **(D)** and *rab7a;rab7bb* **(E)**. Black arrowheads show invaginating luminal front. The final image represents the fully lumenized state of the blood vessels around 44 hpf. **F** Schematic representation of how lumen diameter was measured. The diameter of the vessel was measured perpendicular to vessel axis at 3 positions. The membrane marker *Tg(kdrl:mcherry-CAAX)^s916^* was used as reference to how far the lumen expanded. **G** Violin plot showing lumen diameter in *rab7* mutants. Median is indicated by thick black line (wild-type: N= 8 fish, n= 24 blood vessels; *rab7a*: N= 4, n= 10 (maternal-zygotic homozygous); *rab7ba*: N= 8, n= 22 (maternal-zygotic); *rab7a; rab7ba*: N= 10, n= 31 (maternal-zygotic); *rab7a; rab7bb*: N=7, n= 19 (maternal-zygotic).

### rab7 single and double mutants are capable of transcellular lumen formation and fusion

Since Rab7a appears to associate with the apical membrane during transcellular lumen formation (Fig. 1-2), we then investigated the role of Rab7 in transcellular lumen formation. To do so, we examined the process of anastomosis and focused only on blood vessels that connect and expand their lumen via this process. To differentiate between the two different lumen formation mechanisms (cord hollowing vs transcellular lumen), we imaged the process in *Tg(fli:Pecam1-EGFP)* embryos, which allows us to follow cell-cell junctions and thus identify individual endothelial cells. In transcellular lumen formation, the lumen forms before junctional rearrangements take place and the newly forming luminal connection is surrounded by a single cell only, visualized by a clearly defined junctional ring and the absence of junctions along the endothelial cell body (see Fig 6A). Henceforth, the lumen expands inside a single cell. If the lumen fusion observed in Fig 6A would have been brought about by cellular rearrangements only (cord hollowing), the visualization of a live junctional marker would have revealed junctions running along the entire vessel.

**Figure 6.**
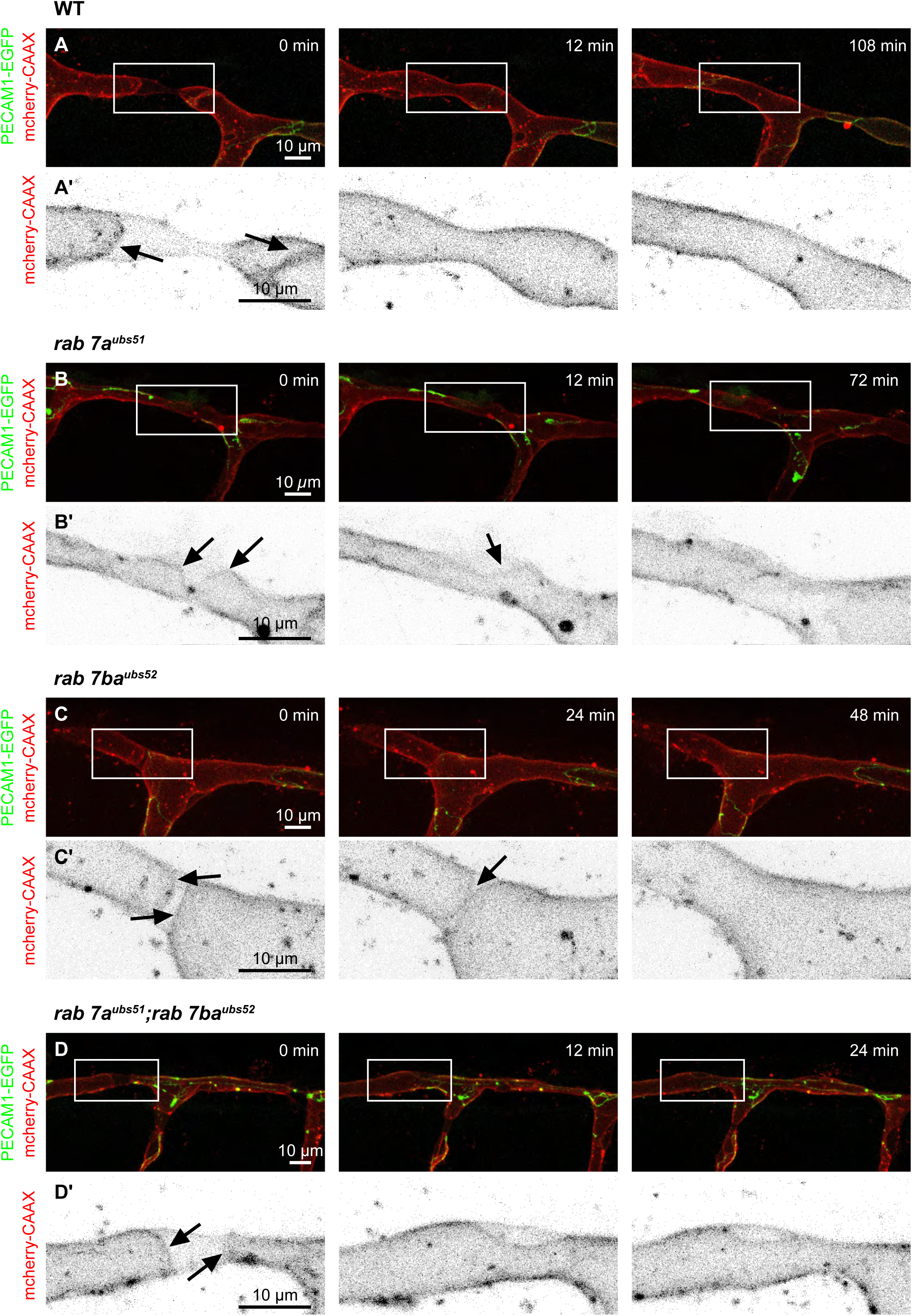
Lumen fusion in *rab7* mutants. **A-D** Stills from high resolution confocal imaging of lumen fusion in the DLAV (anterior to the left) of double transgenic *Tg(kdrl:mcherry-CAAX)^s916^; Tg(fli:Pecam1-EGFP)^ncv27^* wild-type (**A**), maternal-zygotic homozygous *rab7a* double (**B**), maternal-zygotic homozygous *rab7ba* (**C**) and maternal-zygotic double homozygous *rab7a^ubs51^; rab7ba^us521^* (**D**) embryos. **A’-D’** Isolated mcherry-CAAX signal labelling the apical membrane of ROIs from A-D. Arrows indicate the invaginating luminal front.

Time-lapse live imaging of *rab7* mutants expressing both the *Tg(fli:Pecam1-EGFP)* and the *Tg(kdrl:mcherry-CAAX)* marker in endothelial cells showed that transcellular lumen formation was indeed observed in *rab7a* and *rab7ba* single maternal-zygotic homozygous and in *rab7a; rab7ba* double maternal-zygotic homozygous embryos (Fig 6B-D; higher magnification of membrane fusion in Fig 6B’-D’). In these movies, the apical luminal fronts were observed as they grew towards each other and fused upon contact, thereby forming one continuous lumen in a stretch of vessel characterized by the absence of continuous junctions, indicating that this lumen had fused within a single endothelial cell.

### rab7 single and double mutants show differences in late endosomal/lysosomal vesicle size but not in vesicle number

A major function of Rab7 is to control fusion of vesicles in the last step of endosomal trafficking. These late endosomes eventually fuse with lysosomes such that cargo can be degraded. This fusion is orchestrated by Rab7 with the help of its effectors. To investigate whether the trafficking of late endosomes to lysosomes is affected in various *rab7* mutant combinations, the size and number of vesicles positive for the late endosomal/lysosomal marker Lamp2 were measured. The measurement was taken in a defined area around the dorsal-most end of a non-lumenized blood vessel, and all Lamp2-RFP positive vesicles were measured manually (Fig 7 A-C). These measurements showed that vesicle size was significantly reduced in *rab7ba^ubs52^* homozygous embryos as well as in *rab7a ^ubs51^; rab7ba ^ubs52^* double homozygous embryos, when compared to controls (Fig 7A-C). In *rab7a ^ubs51^*, vesicle size was not altered. Lamp1 is sorted into late endosomes and lysosomes from the TGN (Cook et al., 2004) and it has been postulated that *rab7b* has a different function from *rab7a* and plays a role in the shuttling from the TGN to late endosomes (Progida et al., 2010). This might indicate that the observed effect is mostly due to improper trafficking of Lamp2-RFP to late endosomes/lysosomes with the help of Rab7b, and that the effect of Rab7a cannot be studied using the assays we chose. Therefore, a small screen was performed using splice-morpholinos (MOs) against all three isoforms of the zebrafish *rab7* genes, blocking splicing of exon2 (Fig 7G-J). Strikingly, injection of splice-MO in *TgBAC(Lamp2-RFP)* embryos reveals that in embryos injected with MO against *rab7ba*, the Lamp2-RFP signal was lost in the entire embryo compared to standard-MO injected siblings, or to uninjected control embryos. Measurements of vesicles in embryos injected with splice-MO against either *rab7a* or *rab7bb* showed a strong and significant increase of vesicle size compared to standard-MO injected embryos (Fig 7G, I and J). Together, these results indicate that loss of *rab7ba* might play a role in proper localization of Lamp2-RFP. Our findings also indicate that vesicle size in embryos mutant for the two functionally redundant *rab7a* and *rab7bb* is expected to be increased, in line with the MO data, and that, in order to use the Lamp2-RFP signal as a marker, Rab7ba needs to be present.

**Figure 7.**
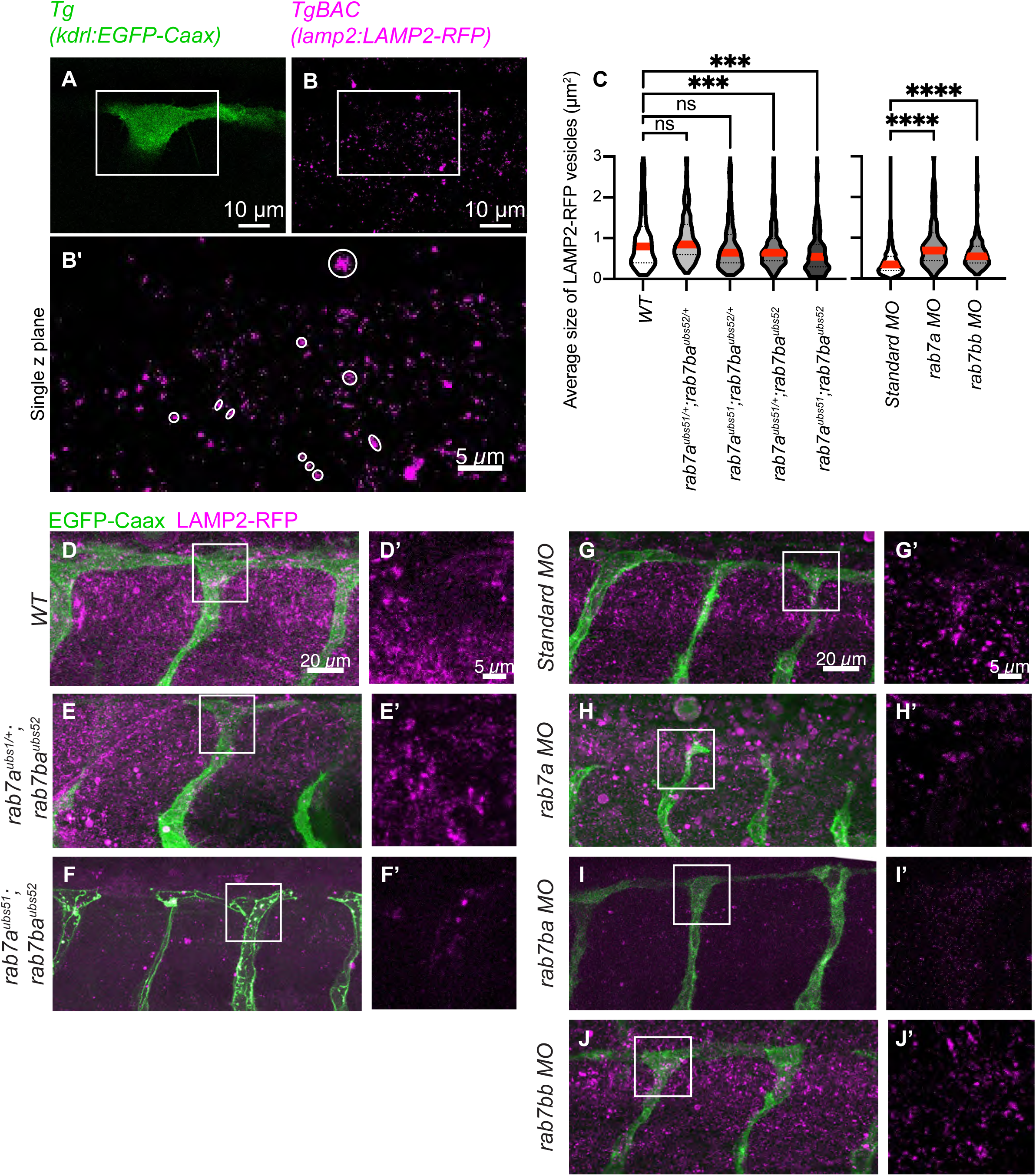
Vesicle analysis in *rab7* loss-of-function. **A-B ’** Representation of measurement of vesicle size. Single z-stack of a non-lumenized tip-cell is chosen using the endothelial marker *kdrl:EGFP-CAAX*. **B’** In a zoom-in window of the ROI in B, every Lamp2-RFP positive signal is measured (using as upper cut off 2 pixels). **G** Violin plots of measured vesicle sizes in *rab7* mutants (wild-type N= 5, n= 283, *rab7a;rab7ba* heterozygous N= 4, n= 347, *rab7a* homozygous *rab7ba* heterozygous N= 5, n=356, rab7a heterozygous; rab7ba homozygous N= 7, n= 539, rab7a;rab7ba homozygous N= 6, n= 260, p***<0.001, red line indicates the median) and in *rab7* morphants (standard-MO N= 8, n= 930, *rab7a*-MO N= 6, n= 129, *rab7bb*-MO N= 6, n=259, p****<0.0001, red line indicates the median). **D-J** Confocal images expressing an endothelial marker and the late endosome marker Lamp2 (double transgenic embryos *Tg(kdrl:EGFP-CAAX);TgBAC(Lamp2-RFP)^pd1117^* or *Tg(fli:Pecam1-EGFP)^ncv27^;TgBAC(Lamp2-RFP)^pd1117^* . **D’-J’** Zoom-in areas indicated in D-J showing isolated Lamp2-RFP signal. **D-F** Images from wild-type or *rab7* mutant embryos. **G-J** Images from embryos injected with a standard morpholino or *rab7* morpholinos.

## Discussion

### The role of rab7bb

Many genes are duplicated in the zebrafish genome when compared to genomes of mammals. This is also the case for *rab7,* for which an additional copy is present in the zebrafish genome when compared to mammals. We find that *rab7ba* and *rab7bb* are more closely related to each other than to the third copy, *rab7a,* and lie in chromosomal regions that share many genes. We also find that the previously unstudied *rab7* gene, *rab7bb,* is expressed in zebrafish at a similar level as *rab7a*. Using targeted mass spectrometry, we show that, at the protein level, Rab7a is the most abundant of the three Rab7 proteins, with almost double the levels of Rab7bb and four to five times the levels of Rab7ba. This is in line with previously acquired transcriptomics data (Lawson et al., 2020). We also show that combined loss of *rab7a* and *rab7bb* increases the severity of all observed phenotypes in single *rab7a* mutants (yolk granules, survival of embryos and increase in lumen diameter). This is not the case in *rab7a; rab7ba* double mutants. This indicates that *rab7bb* alone is sufficient to supplement for *rab7a* function and has thus overlapping functions with *rab7a;* no such redundancy is seen with *rab7ba*. This might also be due to the much lower levels of Rab7ba; further analyses would be required to definitely answer this issue.

### Targeted mass spectrometry of rab7 mutants

To validate our *rab7* mutant lines, we used a targeted mass spectrometry approach. Reference peptides were designed that are either specific for a single proteoform, or are shared between two or three of the different Rab7 proteins. Analysis of maternal-zygotic homozygous mutant embryos of a single *rab7* allele revealed that none of the mutant alleles express significant levels of the equivalent wild-type Rab7. Additionally, the reference peptides would also detect any protein translated and expressed from alternative/cryptic translation start sites. Hence, our results reveal that there are no shortened fragments of Rab7 produced in the respective mutants. Together, these observations strongly indicate that the three mutant alleles we generated represent null alleles. Furthermore, in none of the individual mutants did we find upregulation of any of the other two wild-type Rab7 isoforms at the protein level. This demonstrates that wild-type levels of the other *rab7* genes are sufficient to rescue zebrafish embryos to adulthood.

### rab7 maternal contribution and survival

The analyses of the mutant *rab7* lines we generated revealed that the total loss of Rab7 is lethal, similar to what has been reported in other organisms (Kawamura et al., 2012; Poteryaev et al., 2007). While zebrafish lacking all three copies of *rab7* were identified at 24hpf, these embryos die before reaching adulthood. However, our results also show that the lowest expressed gene, *rab7ba,* is sufficient for 10% of the embryos mutant for the two other alleles (*rab7a; rab7bb*) to survive. We find that the two *rab7* alleles, *rab7a* and *rab7bb*, which were predicted by transcriptomics data to be expressed before maternal-to-zygotic transition, are indeed required for early embryo development. 1-cell stage zebrafish embryos lacking maternal contribution of *rab7a* show a phenotype with enlarged yolk granules, caused most likely by a defect in the deposition and fusion of yolk granules with lysosomes, as has previously been described in *C. elegans* (Poteryaev et al., 2007). This phenotype is more severe when both *rab7a* and *rab7bb* are lost. For *rab7a,* we demonstrate that this phenotype is linked to maternal contribution.

### Rab7 in vesicular trafficking and vascular lumen formation and the different role of Rab7ba

Rab7 co-localises with the membrane marker CAAX and the late endosomal/lysosomal marker Lamp2 at dots, which co-migrate along the expanding apical surface during transcellular lumen formation. Although other members of the endocytic pathway (Rab5c: early endosome; Rab11a: recycling endosome) are present at dots or in proximity to the apical membrane, they did not co-localise with the marker CAAX. Our data suggest that the presence of the late endocytic pathway at apical surfaces might be linked to lumen formation and expansion, similar to what has been proposed for the trachea system in Drosophila (Caviglia et al., 2016, 2017).

Further analysis of the late endosomes/lysosomes using the Lamp2 marker in *rab7* mutants, revealed that in the absence of Rab7ba, endosomal/lysosomal size is reduced. This is independent of whether Rab7a is present or not. Strikingly, the Lamp2-RFP signal is lost in *rab7ba* morphants. It has been previously shown that *rab7a* and *rab7b* exert different functions in vesicular trafficking and *rab7b* was proposed to shuttle newly synthesized hydrolases and lysosomal membrane proteins such as Lamp1 to the late endosome/lysosome from the TGN (Cook et al., 2004; Progida et al., 2010). Our results indicate that in zebrafish, this function is mediated by Rab7ba. We have not been able to analyse the expression of Lamp1 in wildtype or any of the *rab7* mutants, due to the lack of the required tools, such as specific antibodies or transgenic reporter lines. Our data indicate that all Rab7 isoform share some functions but apparently carry out independent functions.

To investigate the potential role of different *rab7* alleles in apical membrane fusion and lumen formation in the zebrafish vasculature, we performed *in vivo* live imaging using markers labelling endothelial membranes (*Tg(kdrl:mcherry-CAAX*)). Vascular development was unaltered in all mutants and mutant combinations analysed, and all blood vessels formed normally and were perfused. However, when we measured lumen diameter in the different mutants, we observed two trends: 1) the lumen was increased in *rab7a* and *rab7a; rab7bb* single and double mutants, respectively, and 2) the lumen was decreased in *rab7ba* and *rab7a; rab7ba* single and double mutants, respectively, meaning that Rab7a and Rab7ba exert different effects on lumen properties. Lumen diameter can be a readout for junctional stability, cell rearrangement and/or blood pressure (Red-Horse and Siekmann, 2019). We did not further investigate how the observed phenotypes were related to Rab7 function.

To further address the role of Rab7 in transcellular lumen formation, we used the apical marker (*Tg(kdrl:mcherry-CAAX*)) together with the endothelial junction marker *Tg(fli:Pecam1-GFP).* We found that in *rab7a* and *rab7ba* single mutants and in *rab7a; rab7ba* double homozygous mutants, transcellular lumen formation and lumen fusion takes place in a manner comparable to wild type embryos.

In order to determine whether the complete absence of Rab7 results in more prominent defects in sprouting angiogenesis and vascular lumen formation, in particular during apical membrane fusion, additional approaches will be required. A conditionally inactivatable allele of *rab7* could be expressed in a triple mutant embryo to rescue lethality; inactivation of this allele in the vasculature, either by genetic means or by protein degradation tools (Harmansa and Affolter, 2018; Yamaguchi et al., 2019) should allow to unravel a potential role of Rab7 in lumen formation *in vivo*. The generation of the required tools, and the validation and analyses of such novel approaches, is beyond the scope of the work described here.

## Material and Methods

### Zebrafish husbandry

Zebrafish were maintained in standard housing conditions according to FELASA guidelines (Aleström et al., 2020). Experiments were performed in accordance with federal guidelines and were approved by the Kantonales Veterinäramt of Kanton Basel-Stadt (1027H, 1014HE2, 1014G). The following zebrafish transgenic lines were used: *Tg(kdrl:mCherry-CAAX)^S916^* (Hogan et al., 2009); *Tg(fli1a:EGFP)^ƴ1^* (Lawson and Weinstein, 2002); *Tg(kdrl:EGFP-CAAX)^ubs47^* (this study), *TgBAC(Lamp2-RFP)^pd1117^* (Rodríguez-Fraticelli et al., 2015), *Tg(fli1:EGFP-Rab7a)^ubs48^* (this study)*, Tg(fli1a:Pecam1a-EGFP)^ncv27^* (Ando et al., 2016), *rab7a^ubs51^* (this study), *rab7ba^ubs52^* (this study), *rab7bb^ubs53^* (this study).

### Constructs

Constructs were cloned using the Multisite Gateway Three-Fragment Vector Construction System (Thermo Fisher Scientific) and destination vectors (ie. pDestTol2CG2) from the Tol2Kit (Kwan et al., 2007).

### fli1:EGFP-rab7a

For generation of the *fli1: EGFP-rab7a* vector, a pDestTol2CG2-heart-gfp with the independent marker cmlc2:EGFP, a fli1 P-5’entry clone (Addgene, Lawson Lab), an EGFP p-middle entry (Addgene, Kwan, Chien lab) and rab7a p-3’entry clone (Clark et al., 2011) were used.

### fli1:EGFP-rab5c

For generation of the *fli1: EGFP-rab5c* vector, a pDestTol2CG2-heart-gfp with the independent marker cmlc2:EGFP, a fli1 P-5’entry clone (Addgene, Lawson Lab), an EGFP p-middle entry (Addgene, Kwan, Chien lab) and rab5c p-3’entry clone (Clark et al., 2011) were used.

### fli1:EGFP-rab11a

For generation of the *fli1: EGFP-rab5c* vector, a pDestTol2CG2-heart-gfp with the independent marker cmlc2:EGFP, a fli1 P-5’entry clone (Addgene, Lawson Lab), an EGFP p-middle entry (Addgene, Kwan, Chien lab) and rab11a p-3’entry clone (Clark et al., 2011) were used.

### kdrl:EGFP-CAAX

For generation of the *kdrl:EGFP-CAAX* vector, a pDestTol2CG2-eye-bfp with the independent marker beta-crystaline:BFP a kdrl P-5’entry clone (Addgene, Santoro Lab), an EGFP-CAAX p-middle entry (Addgene, Kristen Kwan, Chien lab) and a poly-A p-3’entry clone (Addgene, Kristen Kwan, Chien lab) were used.

### Transgenesis

*fli1:EGFP-rab7a and kdrl:EGFP-CAAX* plasmids were injected into one-cell stage embryos together with *tol2* mRNA (30 pg mRNA and 20-40 pg DNA/embryo) as previously described (Kawakami et al., 2000). Upon selection of G0 founders, the F1 generations were maintained as stable transgenic lines (*Tg(fli1:EGFP-Rab7a)^ubs48^*, *Tg(kdrl:EGFP-CAAX)^ubs47^*).

### gRNA synthesis

DNA oligonucleotides encoding gRNAs with invariant adapter sequence were used for each *rab7* gene and were designed using the CHOPCHOP online tool (https://chopchop.cbu.uib.no). For gRNA synthesis, each of the gene specific primers (specific sequence in red; *rab7a* TAATACGACTCACTATAGGGCTCTGACACTATGACATCAGTTTTAGAGCTAGAAATAGCAAG*, rab7ba* TAATACGACTCACTATAGGTTTGAGGAGGACCTTTTTACGTTTTAGAGCTAGAAATAGCAAG *or rab7bb* TAATACGACTCACTATAGGAAGGATGGCTTCTCGTAAGAGTTTTAGAGCTAGAAATAGCAAG) was mixed with the constant oligonucleotide (AAAAGCACCGACTCGGTGCCACTTTTTCAAGTTGATAACGGACTAGCCTTATTTTAACTTGCTATTTCT AGCTCTAAAAC), containing a complementary adapter and a Cas9 recruiting sequence. The resulting DNA was purified by Gel and PCR clean-up Kit (Macherey Nagel) and 0.2 μg of DNA was used for RNA *in vitro* transcription by T7 Megascript Kit (Ambion) according to the manufacturer’s protocol.

### Cas9 protein production

Addgene plasmid pET-28b-Cas9-His was used for Cas9 protein production as previously described (Gagnon et al., 2014). Briefly, the Cas9 protein was expressed in BL21 Rosetta Escherichia coli strain (Novagen) in magic medium at 37 °C for 12 h followed by 24 h at 18 °C. Cells were harvested by centrifugation at 6000rpm for 15 min and stored at 4 °C. The cell pellet was resuspended in 20 mM Tris–HCl buffer (pH 8) containing 0.5 M NaCl and 30 mM imidazole, then ultrasonicated and centrifugated at 140000 rpm at 4 °C for 15 min. The supernatant was loaded on Protino NI-NTA agarose beads equilibrated by the same buffer and incubated for 60min. After 4x washes, protein was eluted with 20mM Tris-HCl buffer (pH 8), containing 0.5 M Imidazole and 0.5 M NaCl, on a column in 1ml stepwise elution. Protein purity was confirmed by SDS-polyacrylamide gel electrophoresis and dialyzed overnight against 20 mM Tris-HCl buffer (pH 8) containing 200 mM KCl and 10mM MgCl_2_ and stored at −80 °C.

### Cas9 protein and gRNA injections

Zebrafish embryos were collected and injected as previously described (Rosen et al., 2009) at one-cell stage using a FemtoJet Injector (Eppendorf) or PV820 injector (WPI) and borosilicate glass needles (outer diameter 1mm, inner diameter 0.5mm, BRAND). For targeted mutagenesis, eggs were injected at one-cell stage with a mixture of gRNA and Cas-9 protein at a 1:1 ratio. Injection mix composition was calculated using the website (https://lmwebr.shinyapps.io/CRISPR_Cas9_mix_calc/) from (Burger et al., 2016). Mutagenesis efficiency was approximately 5% for *rab7a* and approximately 20% for *rab7ba* and *rab7bb*. Germline transmission rate was 30% (3/10) for *rab7a*, 40% (4/10) for *rab7ba* and 50% (3/6) for *rab7bb*.

### Genotyping

For each generated allele, a multiplex four primer PCR was established. The following primers were used for *rab7a*: outer forward primer GGGAAGTCTGTGTGTTTAACAGAAGCCGG, outer reverse primer CCACGCCCCTCTTACTGTTAGTTTGC, mutant specific primer GACATAGTGTCTTTCTTCAAGG, wt specific primer CAGAAGAACTTTCTTCCTTGATGTC. For *rab7ba*: outer forward primer GTGTAAACAGCCACAAGCC, outer reverse primer CACACTGATAGCGTCTATGC, mutant specific primer CCAGAATCCCCTAGGGGAAGCC, wt specific primer CCTCCTCAAAGTGATCATCCTAGG. For *rab7bb*: outer forward primer GTTAGACCCGAACTGCATTTCG, outer reverse primer GAAACCCACATGAACACGG, mutant specific primer GGCTTCTCGTGCTGCTGAAGG, wt specific primer GCAGCACCTTCTTACGAGAAGC. Each PCR results in 3 different bands. A larger non-specific band (outer) and two smaller diagnostic bands (for wildtype or mutant allele). For *rab7a* these are: 645bp (outer), 390bp (wildtype allele), 268bp (mutant allele). For *rab7ba*: 492bp (outer), 223bp (wildtype allele), 297bp (mutant allele). For *rab7bb*: 380bp (outer), 246bp (wildtype allele), 161bp (mutant allele).

### Morpholino Injections

One- to two-cell stage embryos were injected with 4 ng of antisense morpholino oligonucleotide (Gene Tools) targeting the splice donor site of Exon2 of the respective *rab7* gene (*rab7a:* 5′-GTTGATTGCGAGAAACTCACCCGGA-3′; *rab7ba:* 5′-ATGCTGAACAAAACACTTACCCAGA-3′; *rab7bb:* 5′-AAAGCCATCACTTACCCAGAATCCC-3′). All MOs were validated via an RT-PCR assay, in which the absence of Exon2 were validated.

### Image acquisition

Live embryos were selected via their fluorescence signal, anesthetized in E3 with 1x tricaine (0.08%, pH 7, Sigma) and mounted in glass bottom Petri dishes (0.17 mm, MatTek) in 0.7% low-melting-point agarose (Sigma) containing 1x tricaine and 0.003% 1-phenyl-2-thiourea (PTU; Sigma-Aldrich) as previously described (Kotini et al., 2022). For live imaging of lumen invagination, an Olympus SpinD (CSU-W1) spinning disc microscope equipped with a dual camera system and a 60x (NA= 1.5) oil objective was used. Z-stacks were made with a step size of 0.2 μm and frames were acquired every 2-30 sec. For live imaging of vascular development, a Leica SP5 confocal microscopes equipped with a 40×(NA=1.1) water immersion objective was used. Z-stacks were made with a step size of 0.35-0.5μm and frames were acquired every 12-25 min.

### Quantification of vessel diameter

Measurements were done using ImageJ. Blood vessels were measured at three different points along the trunk blood vessels. Measurements were taken perpendicular to the vessel axis at each respective point. At the end an average of all three measurements was plotted.

### Quantification of Lamp2-RFP vesicle size

Vesicle size was analysed manually using ImageJ. An area of interest of 200-200 pixels (1985μm^2^) was selected around the T-shaped tip cell of developing sprouts. Within this area, ROIs were drawn around every Lamp2-RFP positive dot, with an upper cut-off size of 3 pixels. Every single vesicle was plotted individually. As a reference point, the last time frame before the cell was lumenized was used.

### Targeted MS of rab7 proteoforms Sample preparation

Sample preparation was performed using the s-trap protocol (Protifi, NY, US). Here, 10-20 embryos were deyolked using forceps in 1X E3 and immediately stored in an empty Eppendorf tube on ice. Embryos were sonicated using glass beads in Bioruptor in 20μl lysis-buffer (5% SDS, 0.1M triethylammonium bicarbonate (TEAB), 10mM tris (2-carboxyethyl) phosphine, pH 8.5). 20 cycles with 30 seconds on and 30 seconds off were used. Samples were then incubated at 95°C and 300 RPM for 10 min. 1μl of iodoacetamide was added and the samples were incubated in the dark at 25°C for 30min. Not more than 50ug of sample was loaded onto the S-trap column after addition of phosphoric acid to a final concentration of 1.2% and 330μl S-trap buffer (90% Methanol and 10% 1M TEAB, pH 8.5). After a spin down at 4000g for 1min, the column was washed 3 times with S-trap buffer. Afterward the sample was digested using 20μl of digestion buffer and 0.75μg of trypsin. After 1h of incubation at 47°C, the generated peptides were collected. For this, 40μl of S-trap buffer, 40μl of 0.2%formic acid and 35μl of 50% acetonitrile acid were added stepwise to the column followed by centrifugation at 4000g for 1 min in between. Peptides were dried for 1h in a speed vac. Peptides were dissolved in LC buffer (0.1% formic acid in water) and the peptide concentration determined using a SpectroStar nanodrop spectrophotometer (BMG Labtech, Germany) and set to 0.5 ug/uL.

### Targeted Liquid Chromatography-Mass Spectrometry (LC-MS) Analysis

Parallel reaction-monitoring (PRM) assays (Gallien et al., 2012; Peterson et al., 2012) were generated from a mixture of proteotypic heavy reference peptides containing 50 fmol/µL of each (Pan-Rab7 peptides: VIILGDSGVGK, ATIGADFLTK; common Rab7a/Rab7ba peptide: NNIPYFETSAK; Rab7a specific peptides: GADCCVLVFDVTAPNTFK, QETEVELYNEFPEPIK; Rab7ba specific peptide: GADCCVLVYDVTAPTTFK; Rab7bb specific peptides: GADCCVLVYDVTAPNTFK, SNIPYFETSAK, JPT Peptide Technologies GmbH). 2 µL of this standard peptide mix were subjected to LC–MS/MS analysis using a Q Exactive plus Mass Spectrometer fitted with an EASY-nLC 1000 (both Thermo Fisher Scientific) and a custom-made column heater set to 60°C. Peptides were resolved using an EasySpray RP-HPLC column (75μm × 25cm, Thermo Fisher Scientific) and a pre-column setup at a flow rate of 0.2 μL/min. The mass spectrometer was operated in DDA mode. Each MS1 scan was followed by high-collision-dissociation (HCD) of the precursor masses of the imported isolation list and the 20 most abundant precursor ions with dynamic exclusion for 20 seconds. Total cycle time was approximately 1 s. For MS1, 3e6 ions were accumulated in the Orbitrap cell over a maximum time of 50 ms and scanned at a resolution of 70,000 FWHM (at 200 m/z). MS1 triggered MS2 scans were acquired at a target setting of 1e5 ions, a resolution of 17,500 FWHM (at 200 m/z) and a mass isolation window of 1.4 Th. Singly charged ions and ions with unassigned charge state were excluded from triggering MS2 events. The normalized collision energy was set to 27% and one microscan was acquired for each spectrum.

The acquired raw-files were searched using the MaxQuant software (Version 1.6.2.3) against a *Danio rerio* (Zebrafish) database (downloaded from www.uniprot.org on 2021/11/02, in total 46,848 entries) using default parameters except protein, peptide and site FDR, which were set to 1 and Lys8 and Arg10. The search results were imported into Skyline (v21.1.0.278) (MacLean, Tomazela et al. 2010) to build a spectral library and assign the most intense transitions to each peptide. An unscheduled mass isolation list containing all peptide ion masses was exported and imported into the Q Exactive Plus operating software for PRM analysis. Here, peptide samples for PRM analysis were resuspended in 0.1% aqueous formic acid, spiked with the heavy reference peptide mix at a concentration of 2 fmol of heavy reference peptides per 1 µg of total endogenous peptide mass and subjected to LC–MS/MS analysis on the same LC-MS system described above using the following settings: The MS2 resolution of the orbitrap was set to 17,500/35,000 FWHM (at 200 m/z) and the fill time to 50/110ms for heavy/light peptides. AGC target was set to 3e6, the normalized collision energy was set to 27%, ion isolation window was set to 0.4 m/z and the first mass was fixed to 100 m/z. A MS1 scan at 35,000 resolution (FWHM at 200 m/z), AGC target 3e6 and fill time of 50 ms was included in each MS cycle. All raw-files were imported into Skyline software for protein / peptide quantification. To control for sample amount variations during sample preparation, the total ion chromatogram (only comprising precursor ions with two to five charges) of each sample was determined using Progenesis QI software (Nonlinear Dynamics (Waters), Version 2.0) and used for normalization. Normalized ratios were further normalized relative to the control condition and the median ratio among peptides corresponding to one protein was reported

### PRM-MS based Quantification of Rab7 Isoforms

In order to determine how much of the single isoform contributes to the overall abundance of Rab7 protein in zebrafish, measurements with the Pan-Rab7 peptide in *rab7a; rab7ba* double homozygous mutants were taken. In this mutant combination, the remaining signal comes exclusively from *rab7bb* expression. Signal intensity was three times weaker than in wild-type and therefore 31% of the signal detected in the wildtype originates from expression of Rab7bb alone. To determine the amount of Rab7a and Rab7ba in the remaining 69%, measurements with the Pan-Rab7a-Rab7ba peptides, detecting the combined Rab7a/Rab7ba signal, were used. Analysis of *rab7a* and *rab7ba* single mutants show that Rab7a is around 5 times more abundant than Rab7ba. This means that the remaining 69% of wildtype Pan-Rab7 signal is split into 57% Rab7a signal and 12% Rab7ba signal.

### Assembly of phylogenetic tree

To assemble a phylogenetic tree of the *rab7* genes, protein sequences were assembled from ensemble genome browser (https://www.ensembl.org/). The sequences were than listen in a txt file which was uploaded to www.ebi.ac.uk (Madeira et al., 2019). Tree data was then visualized using the phylo.io tool (http://phylo.io) (Robinson et al., 2016). For the species comparison, amino acids of the following genes were used: *Homo sapiens* (human): *rab7A*: ENSG00000075785, *rab7B*: ENSG00000276600; *Mus musculus* (mouse): *rab7a*: ENSMUSG00000079477, *rab7b*: ENSMUSG00000052688*; Drosophila melanogaster* (fruit fly): *rab7*: FBgn0015795; *Danio rerio* (zebrafish): *rab7a:* ENSDARG00000020497, *rab7b:* ENSDARG00000021287, *zgc:100918:* ENSDARG00000087243; *Cyprinus carpio* (common carp): *rab7a*: ENSCCRG00000027229, *rab7b:* ENSCCRG00000044991, *zgc:100918:* ENSCCRG00000016691; *Sinocyclocheilus graham* (golden line barbel): *rab7a:* ENSSGRG00000034717*, rab7b:* ENSSGRG00000031796, *zgc:100918:* ENSSGRG00000020044; *Carassius auratus* (Goldfish): *rab7a:* ENSCARG00000008208, *rab7b:* ENSCARG00000017207, *zgc:100918:* ENSCARG00000004993.

### Transcriptomics data analysis

Transcriptomics data were analysed from two available databases. The Lawson and Li dataset (Lawson et al., 2020) presents transcriptomics data regarding endothelial-specific expression of genes. Developmental stage-specific transcriptomics data was analysed by a currently unpublished RNA-seq dataset which was made publicly available https://www.ebi.ac.uk/gxa/experiments/E-ERAD-475 (thanks to the Busch-Nentwich lab). Data were analysed from both datasets for our genes of interest and visualized using GraphPad Prism.

## Acknowledgements

We thank Michael Bagnat for the *TgBAC(Lamp2-RFP)^pd1117^* zebrafish line and David Dylus for assistance with the phylogenetic tree analysis. We also thank Kumuthini Kulendra for fish care and the Imaging Core Facility of the Biozentrum (University of Basel) for microscopy support. This work has been supported by the Kantons Basel-Stadt and Basel-Land and by grants from the Swiss National Science Foundation (310030_200701 and 310030B_176400) to M.A.

---

**Figure S1.**
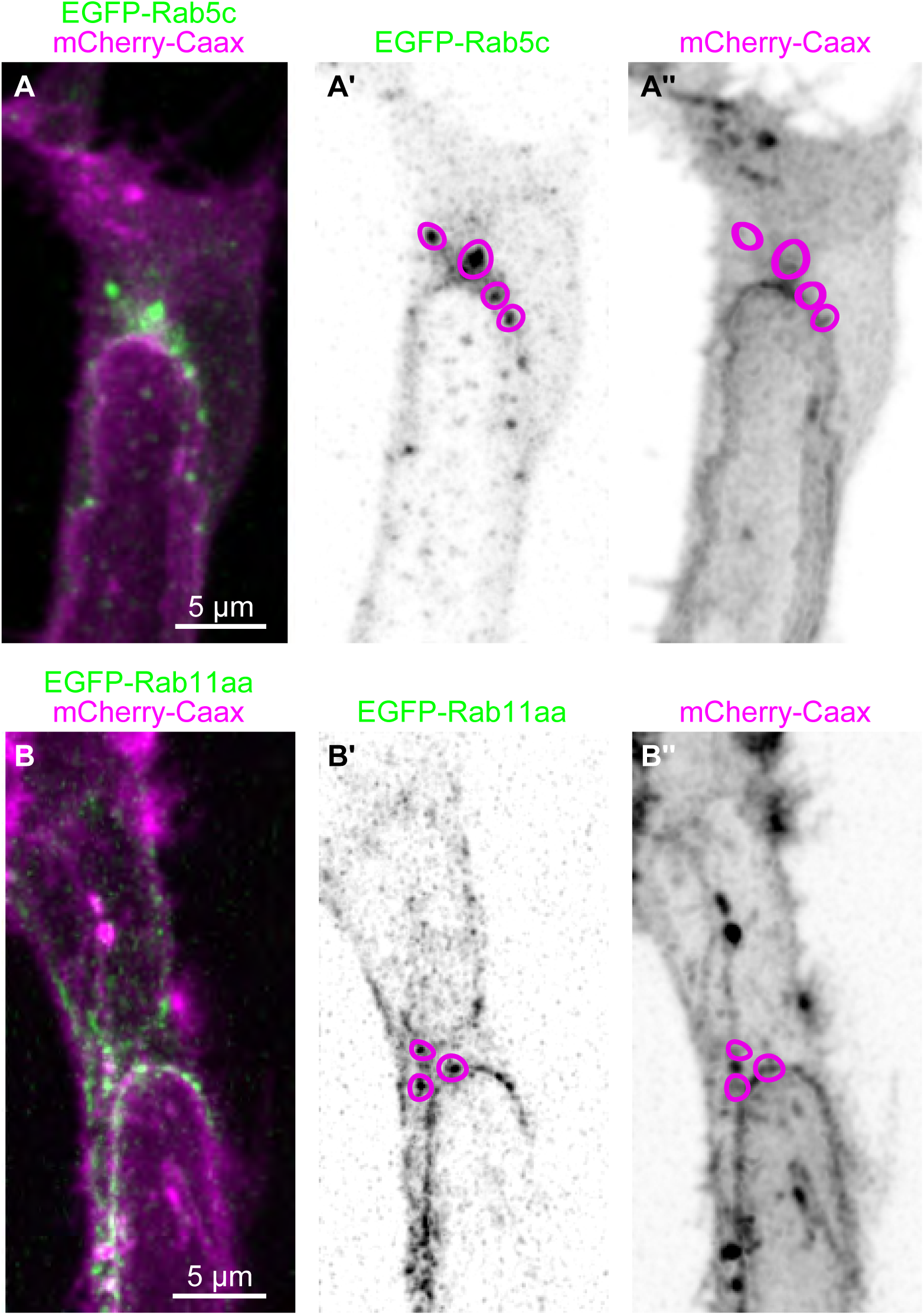
Rab5c and Rab11a localization in endothelial cells. Related to Figure1. **A** Confocal images of a tip cell from a 32hpf transgenic *Tg(kdrl:mCherry-CAAX)^S916^* embryo injected with the plasmid *fli:EGFP-Rab5c* (marker of early endosome). The Rab5c signal (EGFP) does not co-localise with CAAX at dots, but Rab5c dots are found close to the apical surface of the expanding lumen. A’ Inverted contrast of eGFP-Rab5c from A shows Rab5c dots (pink circles). Note that these regions do overlap with the CAAX signal (A’’). A’’ Inverted contrast of mCherry-CAAX from A. **B** Confocal images of a tip cell from a 32hpf transgenic *Tg(kdrl:mCherry-CAAX)^S916^* embryo injected with the plasmid *fli:EGFP-Rab11a* (marker of recycling endosome). The Rab11a signal (EGFP) does not co-localise with CAAX at dots, but Rab11a dots are found in proximity to the apical surface and is also found along the apical membrane of the expanding lumen. A’ Inverted contrast of EGFP-Rab11a from B shows Rab11a dots (pink circles). Note that these regions overlap with CAAX signal (B’’). B’’ Inverted contrast of mCherry-CAAX from B.

**Figure S2.**
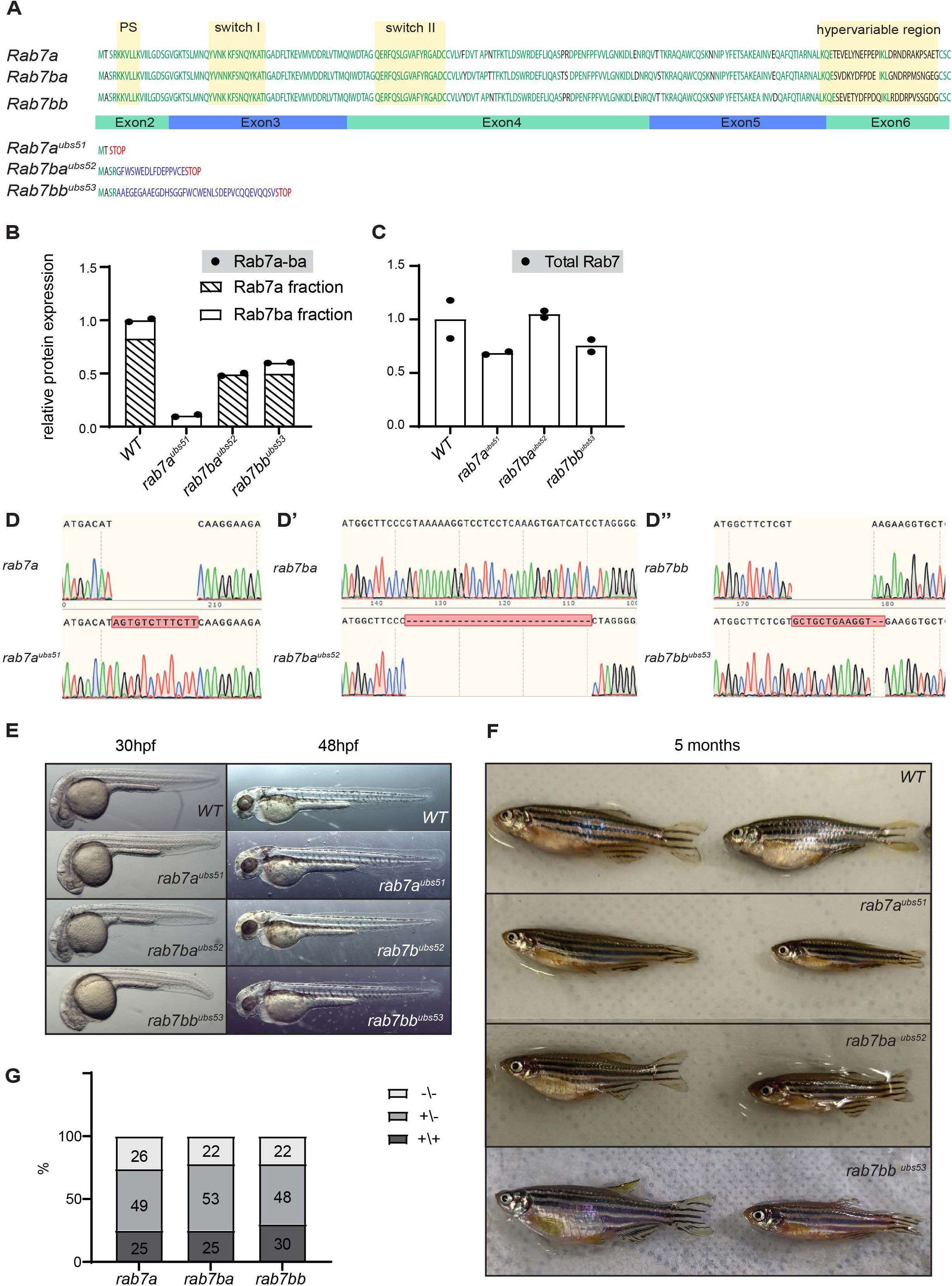
Characterisation of development and survival rates of *rab7* mutants. Related to Figure3. **A** All three *rab7* protein sequences and their predicted mutant sequence. Yellow boxes indicate an important prenylation site responsible for mediating post-translational prenylation of the C-terminal XCXC motif and membrane insertion, and the switch domains important for effector binding after GTP-activation and the hypervariable region responsible for proper membrane recognition. Green letters represent aa that are identical in all three Rab7 isoforms, while black letters represent different aa in Rab7 isoforms. In mutant protein sequences, blue letters are aa encoded out of frame and STOP marks the premature terminating codon. **B-C** Individual value scatter plots of relative protein expression of Rab7ba and the total Rab7 amount. Levels were measured in two different pooled samples of wild-type, *rab7a mat-zyg, rab7ba mat-zyg* and *rab7bb mat-zyg* homozygous embryos. Values were then normalized to total amount of protein measured per sample and to the amount of wild-type sample (n= 2 pools of 20 embryos). **D-D’’** Sequencing results of PCR products of the respective *rab7* loci from 3 months old homozygous fish. **E** Brightfield images of wild-type, *rab7a, rab7ba* and *rab7bb* homozygous mutant embryos at 30 hpf and 48 hpf. **F** Images of wild-type, *rab7a, rab7ba* and *rab7bb* adult fish at 5 months. **G** Percentage of *rab7a, rab7ba* and *rab7bb* mutations found in adult fish from a heterozygous incross of the respective mutant (*rab7a*: N= 3 independent experiments, n= 100 fish; *rab7b*a: N= 3, n= 100; *rab7bb* N= 2, n= 70).

**Figure S3.**
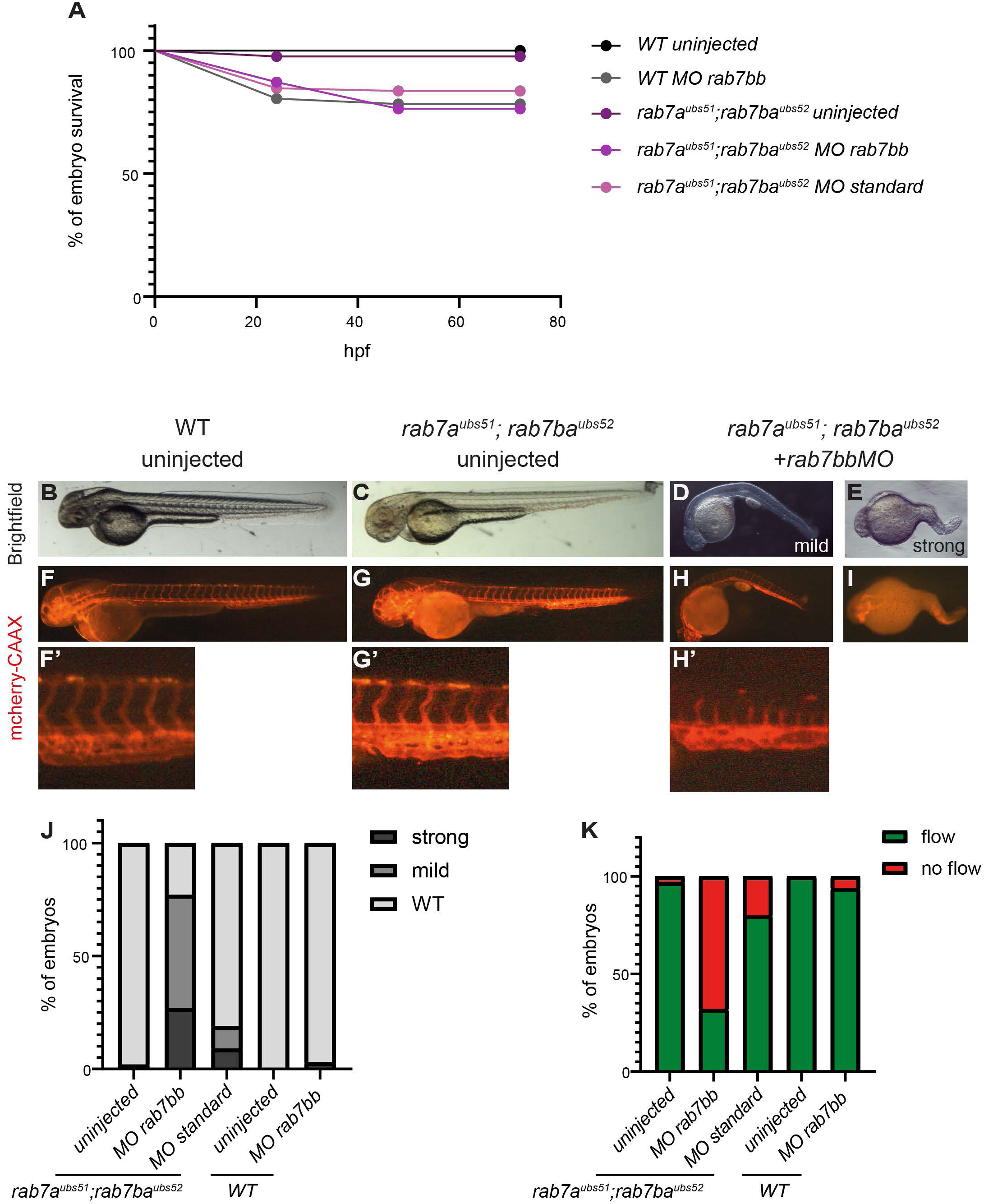
Characterization of development and survival rate of the triple *rab7* loss-of-function mutants. Related to Figure 3 and 4. **A** Percentage of surviving embryos from wild-type, wild-type injected with morpholino against *rab7bb* and *rab7a; rab7ba* mutant incrosses uninjected or injected with control morpholino or *rab7bb* morpholino (n=43-183 embryos per condition). **B-E** Embryo morphology **F-I** and vascular development at 48 hpf of wild-type, double mutant and double mutant injected with the *rab7bb* morpholino (triple loss-of function). **F’-H’** Zoom-in of the boxes in F-H. **J** Developmental defects shown as % of embryos injected with morpholino or uninjected in double mutant or wildtype background (n=42-105 embryos per condition). **K** Presence of blood flow shown as % of embryos injected with morpholino or uninjected in double mutant or wildtype background (n=42-105 embryos per condition).

